# Using a Pharmacokinetic Model to Design and Evaluate an Early ctDNA Biomarker for Response to Targeted Therapy

**DOI:** 10.1101/2025.01.08.632021

**Authors:** Aaron Li

**Affiliations:** School of Mathematics, University of Minnesota, Minneapolis, MN, USA

**Keywords:** birth-death process, cancer, ctDNA, pharmacokinetics, biomarkers

## Abstract

Early prediction of response to therapy or lack thereof can help physicians plan treatment more efficiently. Biomarkers based on circulating tumor DNA (ctDNA) are promising. However, biomarkers beyond direct comparison to baseline have not been thoroughly explored. We develop a model for ctDNA shedding under targeted therapy that incorporates pharmacokinetics. Using a simulated cohort of virtual patients with varied parameters, we define and analyze a biomarker based on ctDNA samples at baseline, 12 hours, and 24 hours after initiation of treatment. The biomarker identified patients who would achieve partial or complete response with high sensitivity and specificity and was able to match the performance of a neural network classifier. Our result highlights the potential of ctDNA as a biomarker and underlines the importance of early ctDNA data collection.

## 1 Introduction

Evaluation of treatment efficacy is an important challenge for cancer treatment. Ineffective treatment can expose patients to unnecessary side effects and delays in finding effective treatments. In the case of non-small cell lung cancer (NSCLC), a wide array of targeted therapy options are available, such as KRAS inhibitors, EGFR inhibitors, BRAF inhibitors, and more [4]. However, the breadth of options can present challenges to physicians trying to design optimal treatment regimens for their patients [1]. Biomarkers that are capable of quickly and accurately detecting response to treatment or lack thereof may help physicians more efficiently decide whether to commit to a given therapy option or rule it out.

Circulating tumor DNA (ctDNA) is a form of cell free DNA that is shed by tumor cells primarily upon apoptosis [9]. Next-gen sequencing has made it possible to collect and sequence ctDNA from non-invasive blood samples as a form of liquid biopsy [6]. ctDNA has shown promise as an indicator of treatment response. In a comprehensive review, Sanz-Garcia et al.[12] examine a variety of proposed ctDNA biomarkers for treatment response, but note that the field is still nascent. Currently, many ctDNA studies [8, 13] collect samples over the course of weeks or months and generally observe that decrease in ctDNA levels from baseline correlates positively with patient outcomes. Parikh et al.[8] hypothesize that earlier ctDNA levels (within two weeks of treatment initiation) may not be predictive of patient response because initiation of therapy can cause acute transient increases in ctDNA. Riediger et al.[11] conducted one of the very few ctDNA studies with daily sampling and observed a sharp transient ctDNA peak immediately after initiation of treatment in a NSCLC patient treated with afatinib, an EGFR inhibitor. These complex ctDNA kinetics may require biomarkers that are more nuanced than decrease from baseline. Mathematical modeling may be able to help define and analyze potential ctDNA biomarkers.

While some mathematical modeling of ctDNA exists, the use of mathematical models in the development of ctDNA biomarkers of treatment response is still under-explored. Avanzini et al.[2] developed a model for ctDNA shedding without treatment and showed that ctDNA sampling may help detect the emergence or reemergence of tumors earlier than imaging. Khan et al.[5] used an ODE model to explore how the presence and quantity of RAS mutations in ctDNA can be used to predict time to treatment failure in colorectal patients treated with cetuximab. Rachman et al.[10] used a spatial model to explore how spatial considerations affect heterogeneity in ctDNA and the tumor population. Li et al.[7] developed a model for ctDNA shedding under targeted therapy, chemotherapy, and radiation treatment and defined biomarkers for treatment response. We now expand upon the targeted therapy model by including pharmacokinetic effects.

In this work, we develop a mechanistic model for tumor and ctDNA kinetics under targeted therapy treatment that incorporates pharmacokinetic dynamics. We use a two-compartment pharmacokinetic model to account for drug absorption and elimination after oral administration. Using NSCLC and afatinib as a motivating example, we parametrize our model with clinical pharmacokinetic data and generate a cohort of 1500 simulated patients. We define a biomarker *V* calculated from ctDNA samples at baseline, 12 hours after treatment initiation, and 24 hours after treatment initiation and show that it can be used to identify patients who will experience partial or complete response with high sensitivity and specificity. We compare the performance of our biomarker with that of a neural network classifier and show that our biomarker is able to match its performance without sacrificing interpretability.

## 2 Model

As in [7], we use a birth-death process with drug-sensitive and drug-resistant subpopulations to model the tumor. We begin with *m*_*s*_ drug-sensitive cells and *m*_*r*_ drug-resistant cells, each of which can divide or die off independently at random exponential rates. The drug sensitive cells have birth and death rates *b*_*s*,1_, *d*_*s*,1_ off drug and *b*_*s*,2_(*t*), *d*_*s*,2_(*t*) on drug, where *b*_*s*,2_(*t*), *d*_*s*,2_(*t*) depend on the pharmacokinetically determined drug-concentration. The resistant cells have constant birth and death rates *b*_*r*_, *d*_*r*_. We assume that sensitive cells and resistant cells are equally viable off drug. That is, *b*_*s*,1_ = *b*_*r*_, *d*_*s*,1_ = *d*_*r*_. Figure 1 shows a schematic of the model.

**Figure 1:**
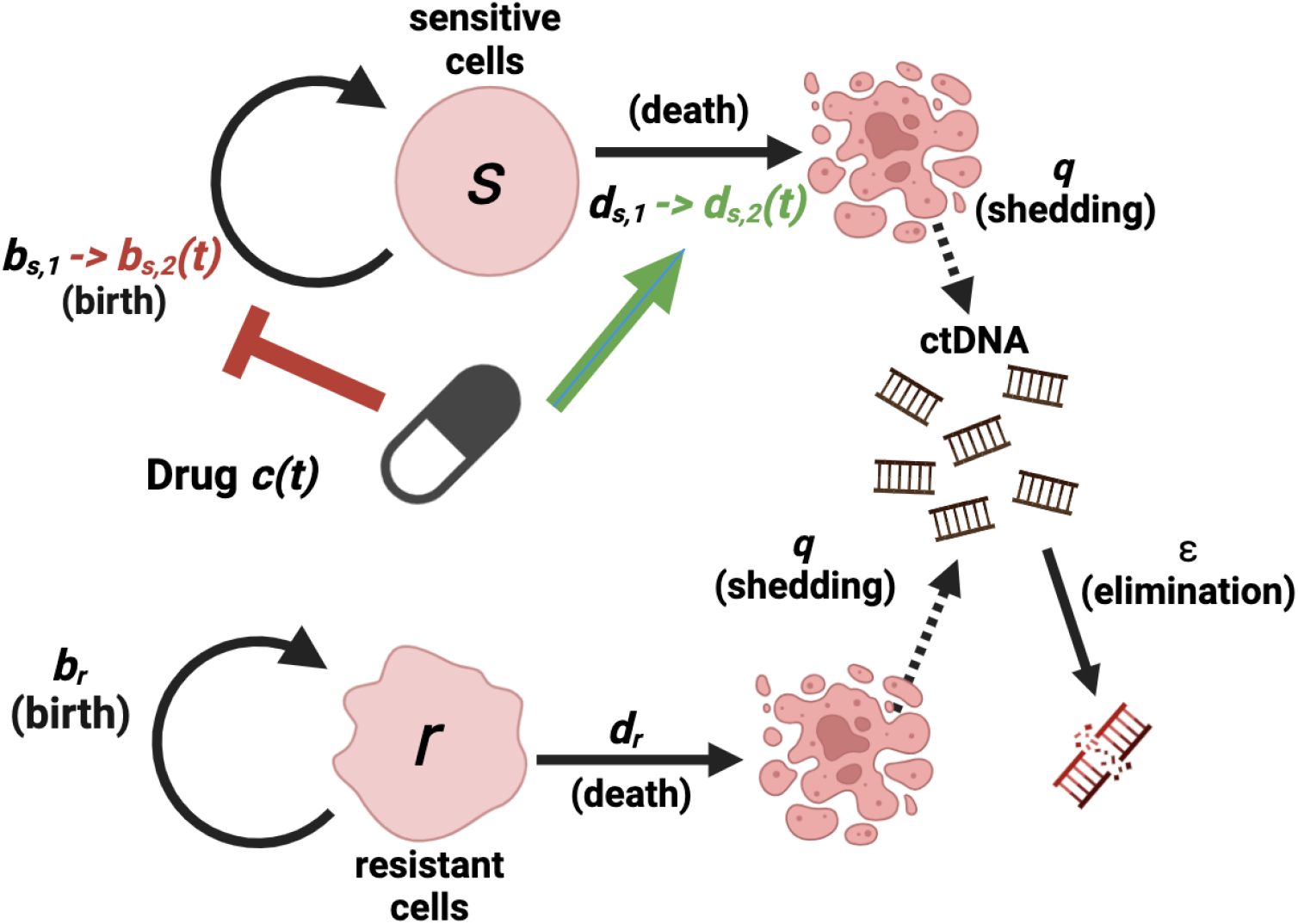
Schematic for ctDNA shedding model. Created in BioRender.

**Figure 2:**
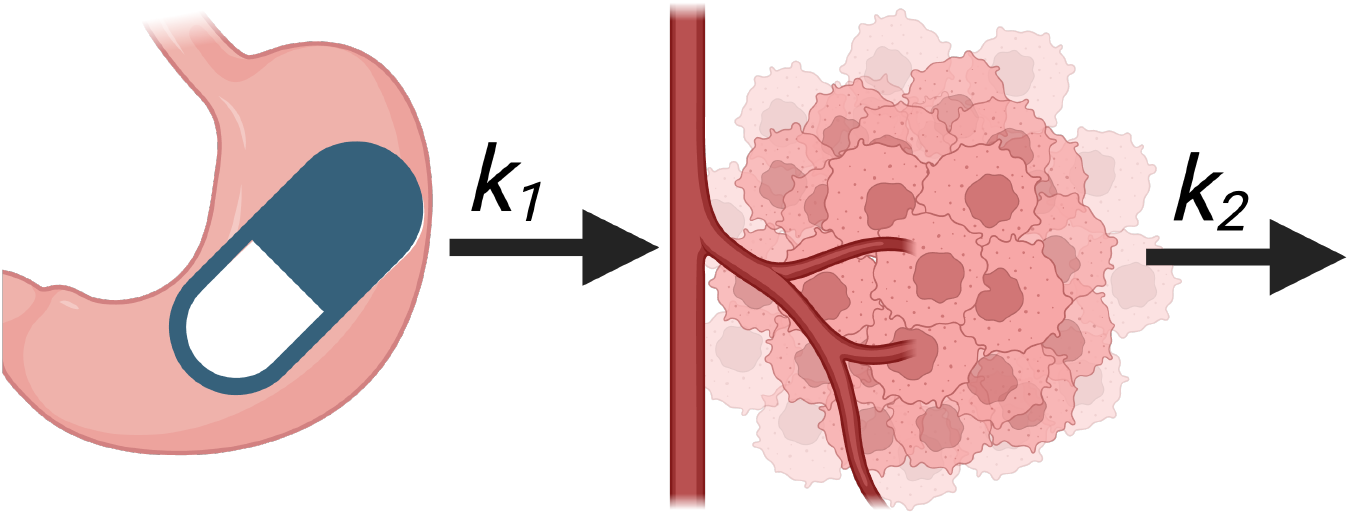
Two-compartment pharmacokinetic model. Created in BioRender.

### 2.1 ctDNA shedding

We base our model for ctDNA shedding on the model proposed by Avanzini et al.[2]. For each cell death, 1 human genome equivalent (hGE) of ctDNA is shed with probability *q*. ctDNA is eliminated from the bloodstream at rate *ε*. We set *ε* = 33/day which corresponds to a ctDNA half life of ∼30 minutes, as has been observed clinically [3]. We assume that both sensitive and resistant cells shed ctDNA at the same rate and we do not assume that ctDNA by resistant cells is distinguishable from ctDNA shed by sensitive cells.

### 2.2 Pharmacokinetics

We use a two-compartment model to describe the pharmacokinetics of orally-administered targeted therapies, such as afatinib. Upon ingestion, the drug is absorbed into the bloodstream from the GI tract at rate *k*_1_. The drug is eliminated from the bloodstream at rate *k*_2_. Let dosage *A* of the drug be administered at times *t*_1_, …, *t*_*n*_, then the total drug concentration in the bloodstream will be

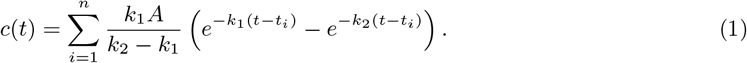

Let *x*_*b*_, *x*_*d*_ be the strength of the drug-effect on the birth and death rates of the drug-sensitive cells. We then define the on-drug birth and death rate of the sensitive cells as follows:

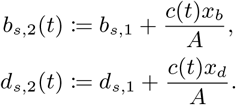

An example simulation is shown in Figure 3. Note the transient peak in ctDNA in the simulation, corroborating the transient peak observed clinically by Riediger et al.[11]

**Figure 3:**
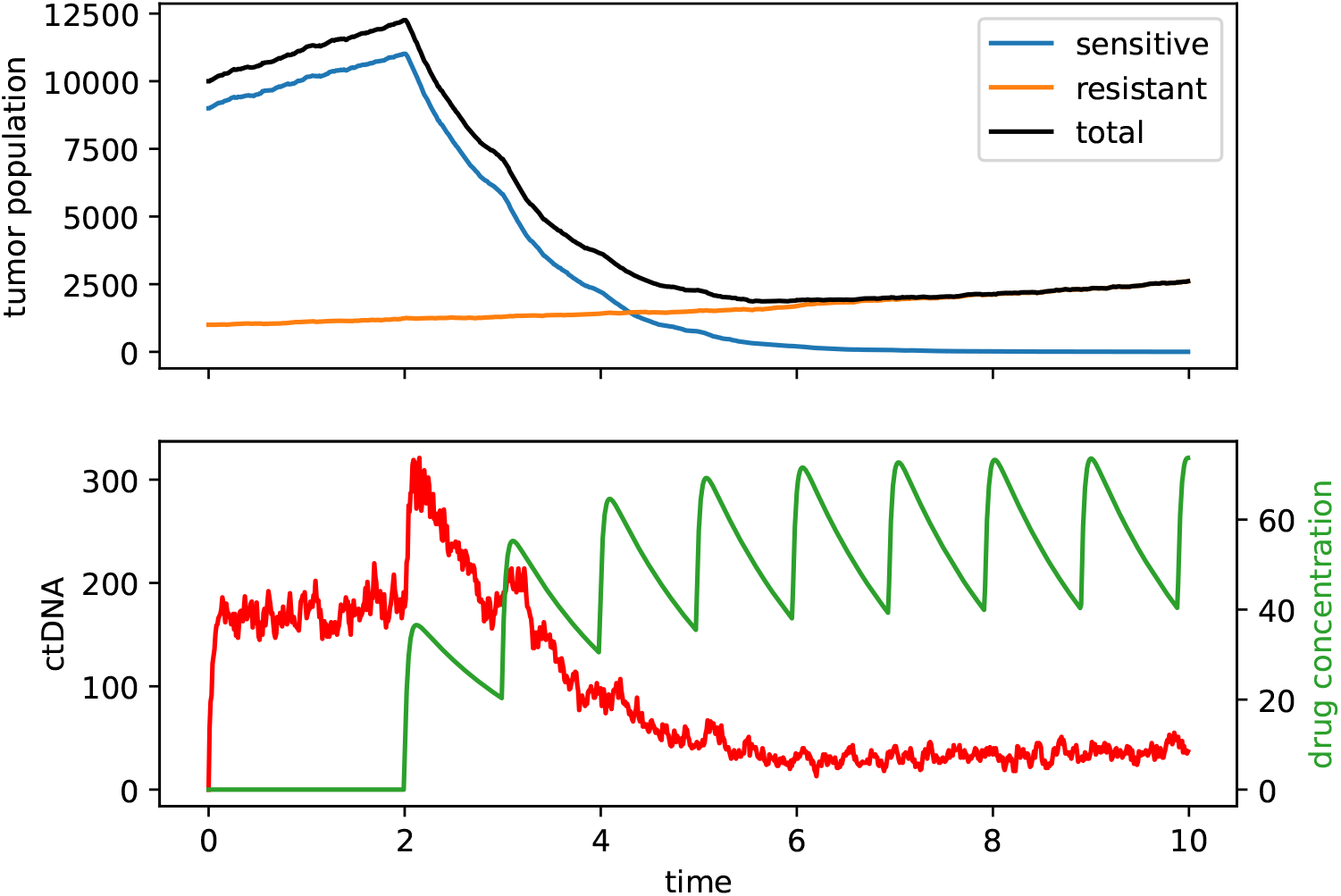
Example simulation with *b*_*s*,1_ = *b*_*r*_ = 1.1/day, *d*_*s*,1_ = *d*_*r*_ = 1/day, *m*_*s*_ = 9000, *m*_*r*_ = 1000, *ε* = 33/day, *q* = 0.5, *k*_1_ = 30, *k*_2_ = 0.7, *A* = 40, *x*_*b*_ = 0, and *x*_*d*_ = 1.

## 3 Simulations

Due to the lack of early ctDNA data, we use a cohort of 1500 simulated afatinib patients to explore the efficacy of ctDNA biomarkers.

### 3.1 Parametrization

For each patient we uniformly randomly select the birth rates *b*_*s*,1_ = *b*_*r*_ in (1.1*/*day, 1.3*/*day), the death rates *d*_*s*,1_ = *d*_*r*_ in (0.8*/*day, 1.0*/*day), the total population *m*_*s*_ + *m*_*r*_ in (2000, 20000), and the proportion of sensitive cells *m*_*s*_*/*(*m*_*s*_ + *m*_*r*_) in (0, 1). We set *q* = 1 and *ε* = 33 for the ctDNA shedding probability and elimination rate.

The simulations start at time *t* = −2 days. At time *t* = 0 days, we simulate the start of daily oral administration of *A* = 40mg afatinib with *x*_*b*_ = 0, *x*_*d*_ = 1. We uniformly randomly select the absorption rate *k*_1_ in (15*/*day, 30*/*day) and the elimination rate *k*_2_ in (0.5*/*day, 1.5*/*day), which correspond to the clinically observed time to max concentration and half life of afatinib [15].

### 3.2 Biomarker definition

For each patient, ctDNA levels are measured at three time points: *t*_0_ = 2, *t*_1_ = 2.5, *t*_2_ = 3. The first measurement serves as a baseline reading and the second and third measurements occur at 12 and 24 hours after initiation of treatment. Let *C*(*t*) represent the ctDNA concentration at time *t*. Then we define a biomarker

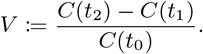

Intuitively, *V* represents the change from *t*_1_ to *t*_2_, normalized by the baseline measurement at *t*_0_. Biomarkers that examine decrease in ctDNA relative to baseline may be limited in predictive power, as noted by Parikh et al. [8], because responsive patients may exhibit sharp transient peaks in ctDNA levels immediately after initiation of treatment [11]. By examining the change from *t*_1_ to *t*_2_ instead of examining change from baseline, our biomarker is able to capture features of ctDNA kinetics without being encumbered by the initial increase in ctDNA levels.

## 4 Results

In order to evaluate the biomarker, we examine its ability to identify patients who will or will not achieve partial or complete response (PR). RECIST 1.1 defines partial response as a decrease in tumor burden of at least 30% [14]. Figure 4 shows a receiver operating characteristic curve for *V* when performing a binary classification of patients who will achieve at least partial response or not. The area under the curve (AUC) produced by *V* is 0.93. At the optimal threshold of *V <* −0.23, partial or complete response was identified with 89% sensitivity and 88% specificity.

**Figure 4:**
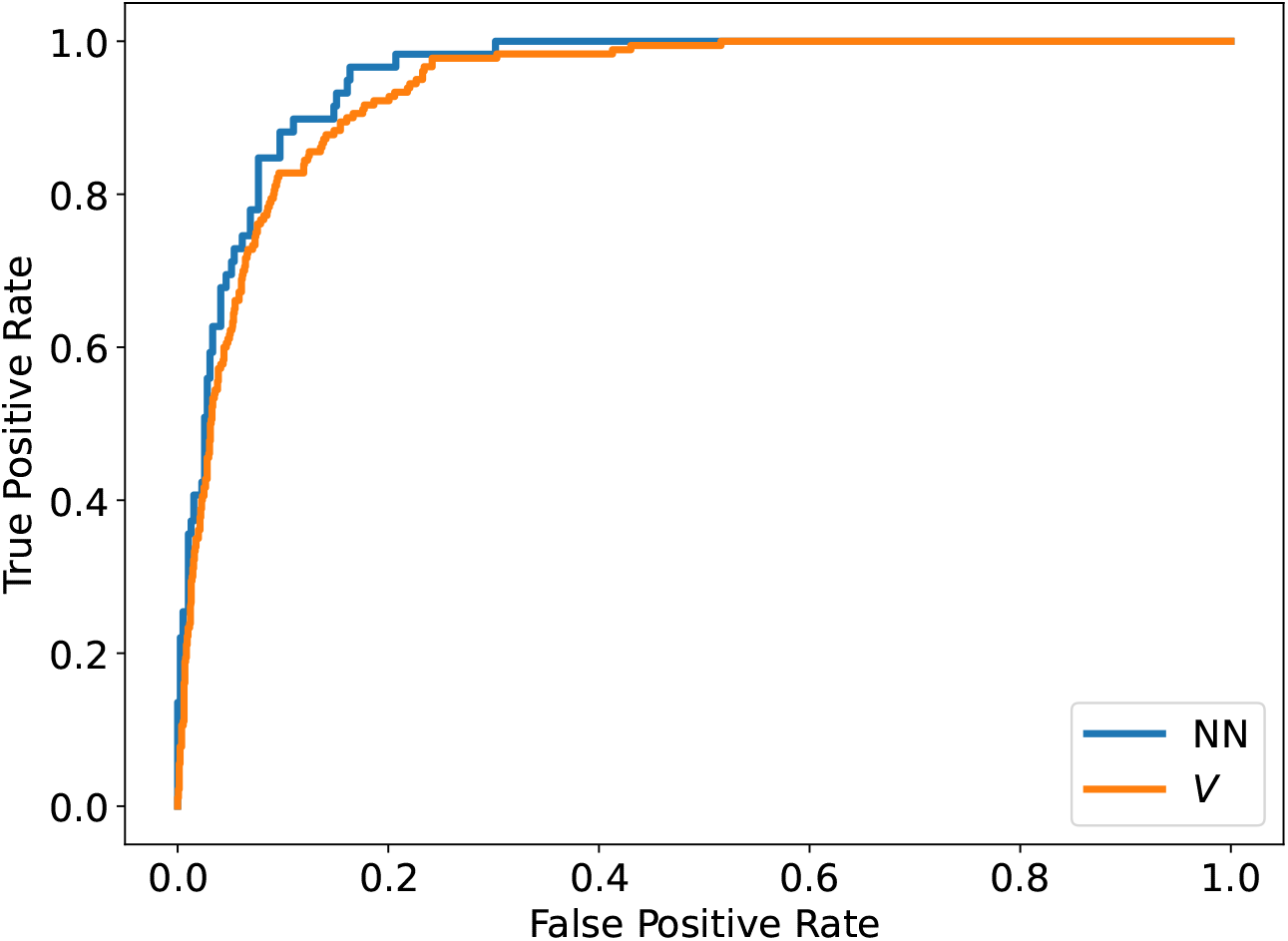
Receiver operating characteristic curves for detection of partial or complete response in our simulated cohort of 1500 patients. The biomarker *V* produced an area under the curve (AUC) of 0.93 while the neural network classifier produced an AUC of 0.95.

### 4.1 Validation against neural network

In order to validate the performance of our biomarker *V*, we compare it to the performance of a neural network classifier. The classifier is a three-layer neural network implemented in PyTorch and was trained on 70% of the simulated cohort for 300 epochs. Given the three ctDNA samples for each patient, the classifier outputs a binary classification of whether the patient will achieve at least partial response or not. When tested on the remaining 30% of the cohort, the neural network classifier produced an AUC of 0.95. Thus, our biomarker *V* is able to closely match the performance of the neural network as shown in Figure 4.

## 5 Discussion

We developed a model of ctDNA shedding under targeted therapy that incorporates a two compartment pharmacokinetic system. Using this model and randomized parameters based on clinical pharmacokinetics of afatinib, we simulated a cohort of 1500 patients. We defined a biomarker *V* = (*C*(*t*_2_)−*C*(*t*_1_))*/C*(*t*_0_) in terms of ctDNA measurements at baseline, 12 hours, and 24 hours after treatment initiation. We found that *V* successfully differentiated patients who would go on to achieve partial or complete response from patients who would not experience reduction in tumor size. The ROC curve of *V* had area under the curve 0.93 (Figure 4) and the threshold of *V <* −0.23 identified responders with 89% sensitivity and 88% specificity. Our biomarker compares favorably to existing ctDNA biomarkers not only in terms of accuracy, but also in terms of earliness of prediction. For example Parikh et al.[8] found that change in ctDNA from baseline to week four of targeted therapy treatment predicted clinical benefit with 90% specificity and 60% sensitivity. Our biomarker would only require sampling over the course of a single day, rather than weeks or months, providing an immediate prediction of response.

To assess whether our biomarker definition was optimal, we trained a neural network classifier on the same data. The neural network did not significantly exceed the performance of our biomarker, suggesting that our biomarker is not missing important hidden features in the data. Moreover, our biomarker is interpretable and intuitive and does not rely on black box calculations unlike the neural network classifier.

While our biomarker analysis is based on a simulated patient cohort, we hope that our results will inform the development of early ctDNA biomarkers and encourage experimentalists to collect more early ctDNA data. Early prediction of a drug’s success or failure will help physicians formulate treatment strategies and allow patients to avoid unnecessary side effects. The limitations to this work also present opportunities for future exploration. Our model does not incorporate any analysis of spatial effects on cell growth, ctDNA shedding, or drug diffusion throughout the tumor. We also do not examine combination treatments or alternate drug schedules. Future work could involve extending the model to such scenarios.

## Acknowledgements

I would like to thank my advisor, Dr. Jasmine Foo for her help and feedback on this project.

